# Long Period Modeling SARS-CoV-2 Infection of in Vitro Cultured Polarized Human Airway Epithelium

**DOI:** 10.1101/2020.08.27.271130

**Authors:** Siyuan Hao, Kang Ning, Cagla Aksu Kuz, Kai Vorhies, Ziying Yan, Jianming Qiu

**Author notes:** **Corresponding author** Jianming Qiu MSN 3029, 3901 Rainbow Blvd. Kansas City, KS 66160 Or Ziying Yan 1-111 BSB, 51 Newton Road Iowa City, IA 52242 Version: 8-28-20d.

## Abstract

Severe acute respiratory syndrome coronavirus 2 (SARS-CoV-2) replicates throughout human airways. The polarized human airway epithelium (HAE) cultured at an airway-liquid interface (HAE-ALI) is an in vitro model mimicking the in vivo human mucociliary airway epithelium and supports the replication of SARS-CoV-2. However, previous studies only characterized short-period SARS-CoV-2 infection in HAE. In this study, continuously monitoring the SARS-CoV-2 infection in HAE-ALI cultures for a long period of up to 51 days revealed that SARS-CoV-2 infection was long lasting with recurrent replication peaks appearing between an interval of approximately 7-10 days, which was consistent in all the tested HAE-ALI cultures derived from 4 lung bronchi of independent donors. We also identified that SARS-CoV-2 does not infect HAE from the basolateral side, and the dominant SARS-CoV-2 permissive epithelial cells are ciliated cells and goblet cells, whereas virus replication in basal cells and club cells was not detectable. Notably, virus infection immediately damaged the HAE, which is demonstrated by dispersed Zonula occludens-1 (ZO-1) expression without clear tight junctions and partial loss of cilia. Importantly, we identified that SARS-CoV-2 productive infection of HAE requires a high viral load of 2.5 × 10^5^ virions per cm^2^ of epithelium. Thus, our studies highlight the importance of a high viral load and that epithelial renewal initiates and maintains a recurrent infection of HAE with SARS-CoV-2.

## Introduction

Coronavirus disease 2019 (COVID-19), an acute respiratory tract infection that emerged in late 2019, is caused by a novel coronavirus, severe acute respiratory syndrome coronavirus 2 (SARS-CoV-2) ^1–5^. It is an enveloped, positive-sense, single-stranded RNA virus, and belongs to the genus *Betacoronavirus* of the family *Coronaviridae* ^1–3, 6^. The clinical syndrome of COVID-19 is characterized with a varying degree of severity, ranging from a mild upper respiratory illness ^7^ to severe interstitial pneumonia and acute respiratory distress syndrome (ARDS), a life-threatening lung injury that allows fluid to leak into the lung ^5, 8–10^. Compared to ~34% fatality of the middle east respiratory syndrome (MERS) and ~10% of the severe acute respiratory syndrome (SARS) ^11^, COVID-19 has a lower fatality rate, ranging from 0.7% to 5.7% in the United States ^12^, however, it spreads more efficiently than SARS and MERS ^13, 14^, making it difficult to contain. COVID-19 has become a pandemic ^15^, which has led to over 24.5 million confirmed cases and >834,000 fatalities worldwide as of August 27, 2020.

While SARS-CoV-2 viral RNA can be detected in nasal swabs, nasopharyngeal aspirates, and bronchoalveolar lavage fluids throughout the airways ^1–3, 16, 17^, how the virus infects epithelial cells at different levels of the respiratory tree and the underlying pathogenesis remain unclear. Thus, a comprehensive understanding of how SARS-CoV-2 replicates and causes pathogenesis in its native host, the epithelial cells lining different levels of the airways, is essential to devising therapeutic and prevention strategies to counteract COVID-19. Primary human nasal, trachea, and bronchial epithelial cells can be cultured and differentiated at an air-liquid interface (ALI), forming a pseudostratified mucociliary airway epithelium that is composed of ciliated cells, goblet cells, club cells and basal cells with an arrangement closely reflective of an in vivo cellular organization ^18, 19^. This in vitro model of human airway epithelium (HAE) cultured at an ALI (HAE-ALI) closely recapitulates many important characteristics of respiratory virus-host cell interactions seen in the infected upper and lower airways in vivo, and has been used to study many human respiratory viruses ^20–29^, including SARS-CoV ^30, 31^. Primary HAE-ALI can be infected by SARS-CoV-2 ^32, 33^ resulting in epithelial damage ^34^, and they can be used for virus isolation ^1, 16^. Notably, differentiation at an ALI results in a drastic increase in expression and the polar presentation of the viral receptor angiotensin-converting enzyme 2 (ACE2) ^2, 35^ on the apical membrane ^30, 31^. Thus, the differentiation/polarization of the epithelial cells is an optimal cell culture model to study SARS-CoV-2 infection in vitro.

In this study, we generated HAE-ALI cultures directly from primary bronchial epithelial cells without propagation prior to differentiation at an ALI and we used these cultures to model SARS-CoV-2 infection. Studies were carried out for a long period of up to 51 days and focused on the viral replication kinetics, the dose dependency of viral infection, epithelial damage and the permissive subpopulation of the epithelial cell types. The polar infection of SARS-CoV-2 in HAE-ALI was confirmed. While SARS-CoV-2 efficiently infected HAE-ALI through the apical side at a viral load as low as a multiplicity of infection (MOI) of 0.002 plaque forming units (pfu) per cell, at an MOI lower than this threshold the viral replication was undetectable. Notably, SARS-CoV-2 infection presented as an enduring infection in HAE-ALI with recurrent peaks of virus released from the infected ciliated and goblet cells while the airway basal cells and club cells are non-permissive.

## Results

### SARS-CoV-2 infection of human airway epithelia presents a long-lasting infection and causes epithelial damage

SARS-CoV-2 primarily infects human airway epithelial cells of the respiratory tract and lung of COVID-19 patients ^36, 37^. SARS-CoV-2 infections in the in vitro model of well-differentiated HAE-ALI or as organoids have been reported ^34, 36, 38, 39^. However, these studies were focused on short-term virus replication and cytopathic effects as they were carried out in a timeframe of less than one-week post-infection. Since airway epithelia capably repair, regenerate, and remodel themselves ^40^, we hypothesized that a long-term monitoring of SARS-CoV-2 infection in HAE-ALI might reveal unknown important features that were missed in prior studies.

To this end, we first chose two HAE-ALI cultures, B4-20 and B9-20 (HAE-ALI^B4-20^ and HAE-ALI^B9-20^), which were respectively generated from primary bronchial epithelial cells freshly isolated from two donors. The initial study was performed with the infection of SARS-CoV-2 at an MOI of 2 pfu/cell. We collected the apical washes on a daily base for continued monitoring of the virus replication through titration for infectious virions with plaque assay in Vero-E6 cells. We also periodically performed confocal microscopy analysis of the infected HAE with immunofluorescent assays. As expected, we observed rapid virus release from the infected HAE-ALI cultures, which reached a peak of 9 × 10^5^ pfu/ml at 2 days post-infection (dpi). The apical virus release from HAE-ALI^B4-20^ remained at the peak for 3 days, then decreased to a level less than 800 pfu/ml at 7 dpi (**Fig. 1**, B4-20). The infection of HAE-ALI^B9-20^ presented a similar trend with the peak at 7.5 × 10^5^ pfu/ml from 2-6 dpi, which dropped to 3 × 10^3^ pfu/ml at 9 dpi (**Fig. 1**, B9-20). However, at later time points, continued study revealed a virus release kinetics with at least two peaks during the course of 3 weeks from both infections. In the infection of HAE-ALI^B4-20^, the virus release started to increase again from 8 dpi and reached a peak of 7.6 × 10^5^ pfu/ml at 12 dpi. The apical release of virus then dropped to 3 × 10^4^ pfu/ml at 14 dpi, followed by another peak of 7 × 10^5^ pfu/ml at 17 dpi. (**Fig. 1**, B4-20). Notably, while the decrease of the virus released from the 1^st^ peak of infected HAE-ALI^B9-20^ lagged behind that of the infected HAE-ALI^B4-20^ by 2 days, it demonstrated the second peak at 11-13 dpi (**Fig. 1**, B9-20), which contained viral shedding at 7.6 × 10^5^ pfu/ml. We reasoned that this was due to the donor variation, which affects the extent of differentiation and the ratio of the subpopulation of epithelial cell types, but not the properties permissive to SARS-CoV-2 infection. To address this, we infected the HAE-ALI ^B3-20^, which was derived from another donor, with a 10-fold reduced virus inoculum (MOI of 0.2 pfu/cell). We observed similar replication kinetics with two virus release peaks (**SFig. 1A**). Of note, even though a reduced MOI was applied to HAE-ALI ^B3-20^, we recorded a higher viral shedding in the 1^st^ peak (4 × 10^6^ pfu/ml) than that from HAE-ALI^B4-20^ and HAE-ALI^B9-20^, while those in the 2^nd^ peaks of the infections in the three cultures were approximately at a similar level (5 × 10^5^ pfu/ml). SARS-CoV-2 infection of HAE-ALI^B3-20^ significantly reduced the TEER values starting at 1 dpi (**SFig. 1B**), caused dispersed ZO-1 expression and loss the cilia (**SFig. 1C&D)**, which will be further discussed below.

**Figure 1.**
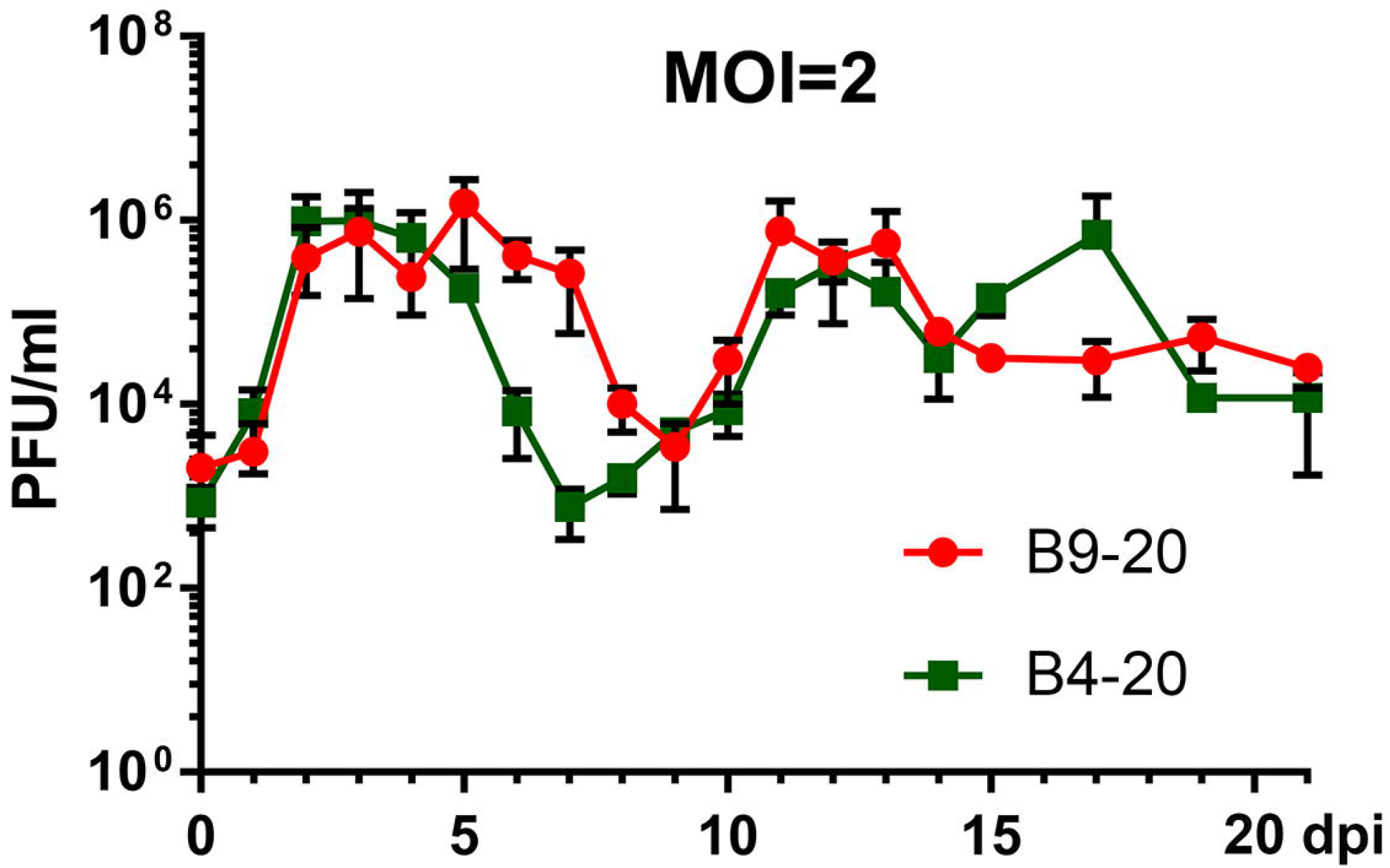
SARS-CoV-2 infection of primary human bronchial airway epithelium (HAE) is persistent. HAE-ALI^B4-20^ and HAE-ALI^B9-20^ cultures were infected with SARS-CoV-2 at an MOI of 2 from the apical side. At the indicated days post-infection (dpi), the apical surface was washed with 100 μl of D-PBS to collect the released virus. Plaque forming units (pfu) were determined (y-axis) and plotted to the dpi. Value represents the mean +/− standard deviations.

SARS-CoV2-infection in HAE-ALI was visualized by immunostaining for the expression of viral nucleocapsid protein (NP) in the infected cells. The analyses revealed the relative increases of NP-positive (NP+) cells, aligned roughly with the apical virus release kinetics (**Fig. 2** and **SFig. 2**, NP). Of note, infected HAE-ALI showed a poor staining of zonula occludens (ZO)-1, which started at 1 dpi and remained throughout the infection, indicating rapid epithelial damage caused by the infection as the tight junctions of the epithelia were destroyed (**Fig. 2A**, **SFig. 1C** and **SFig. 2A**, ZO-1). The Infected HAE-ALI also showed a partial loss of the cilia, indicated by immunostaining with anti-β-tubulin IV, which also started at 1 dpi and remained at a similar level throughout the course of infection (**Fig. 2B**, **SFig. 1D**, and **SFig. 2B**, β-tubulin IV). To observe the infected epithelia in a greater detail, we performed Z-stacked imaging of the infected HAE-ALI^B9-20^ at 15 dpi. The images showed a percentage of ~10% NP+ cells, broken tight junctions (**Fig. 3A**, SARS-CoV2/ZO-1), and an approximate loss of half of the cilia (**Fig. 3B**, SARS-CoV2/Tubulin), compared with the mock-infected HAE (**Fig. 3B**, Mock). Of note, most of the NP+ cells remained β-tubulin IV stained, suggesting that ciliated cells represent the major cell type in HAE permissive to SARS-CoV-2. We also noticed that there were less cells present in the area where NP1 staining were positive, compared to the mock infection, as determined by the number of the nuclei in the imaged area, indicating cell loss (death) of the infected epithelia (**Fig. 3,** DAPI, SARS-CoV-2 vs Mock).

**Figure 2.**
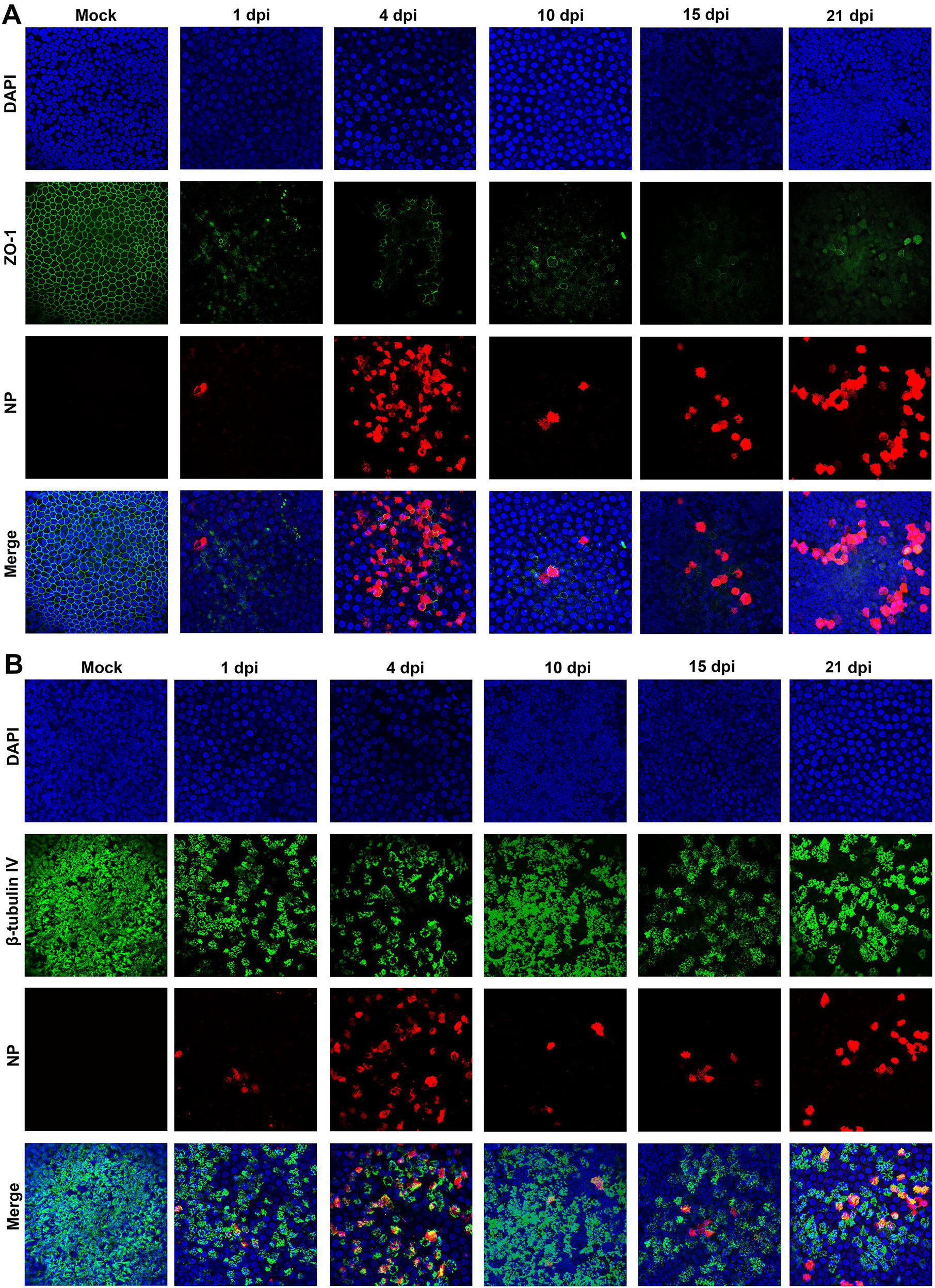
Immunofluorescence analysis of SARS-CoV-2 infected primary bronchial HAE-ALI over a course of 21 days. Mock- and SARS-CoV-2-infected HAE-ALI^B4-20^ cultures were co-stained with anti-NP and anti-ZO-1 antibodies (**A**), or co-stained with anti-NP and anti-β-tubulin IV antibodies (**B**). Confocal images were taken at a magnification of x 40 on the indicated days post-infection (dpi). Nuclei were stained with DAPI (blue).

**Figure 3.**
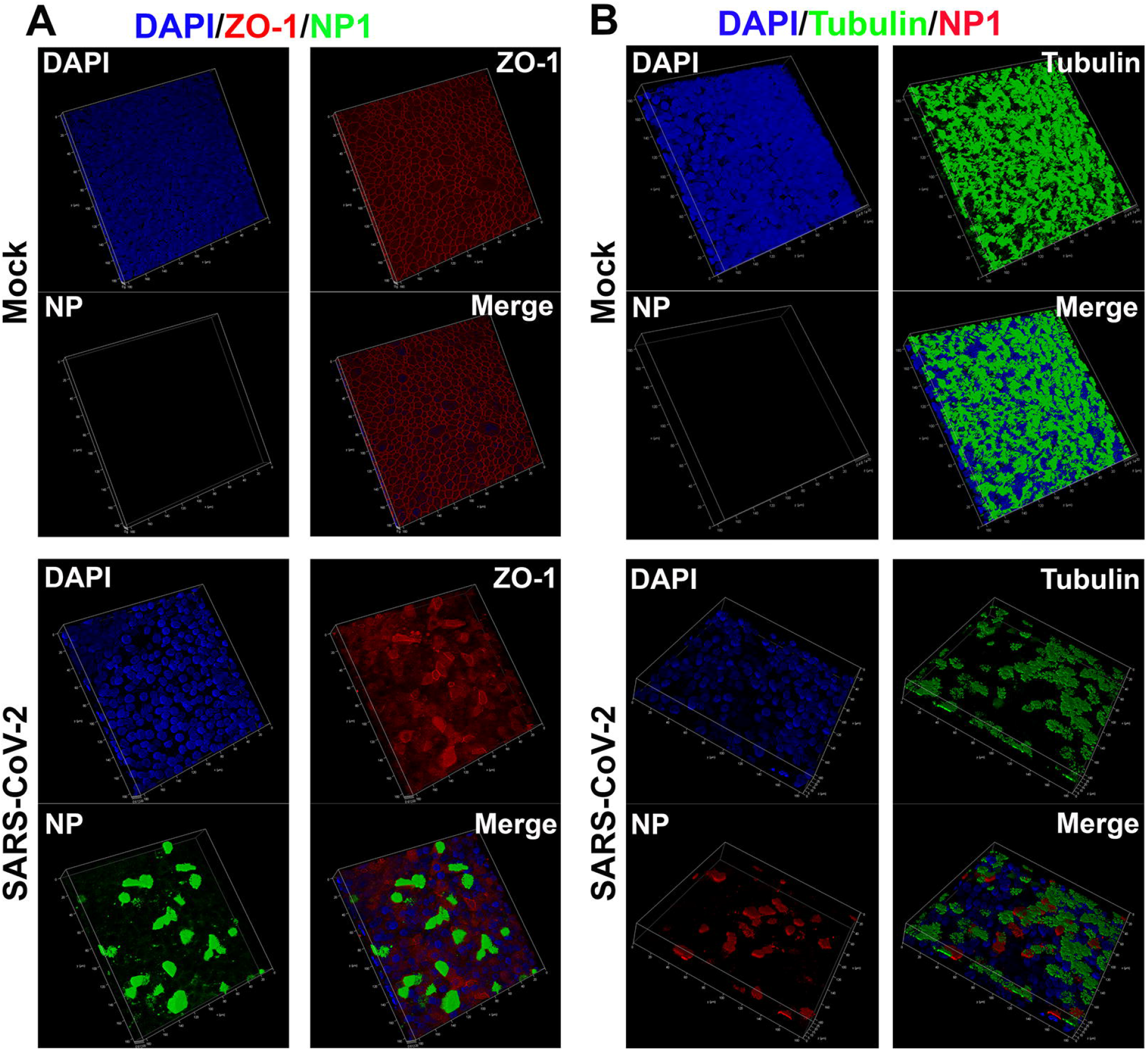
Three-dimensional (z-stack) imaging of SARS-CoV-2 infected primary bronchial HAE-ALI. Mock- and SARS-CoV-2-infected HAE-ALI^B9-20^ cultures at 15 dpi were co-stained with anti-NP and anti-ZO-1 antibodies (**A**), or with anti-NP and anti-β-tubulin IV antibodies (B), or co-stained anti-NP and anti-ZO-1 antibodies (**B**). A set of confocal images were taken at a magnification of x 40 from the stained pierce of epithelium at a distance of the Z value (μm), shown in each image, from the objective (z-axis) and reconstituted as a 3-dimensional (z-stack) image as shown in each channel of fluorescence. Nuclei were stained with DAPI (blue).

Taken all together, these results demonstrated that SARS-CoV-2 infection of HAE-ALI represents a long-lasting process with multiple peaks of virus infections (apical virus release and NP-expressing cells), and that the infection deteriorates two hallmarks of the airway epithelia: the tight junctions and ciliary expression.

### SARS-CoV-2 infection of HAE presents multiple peaks and requires a high viral load

To further examine the recurrent peaks of virus release from the infections and the barrier dysfunction of SARS-CoV-2 infected HAE, we performed a longer monitoring period of 31 days for the infection of HAE-ALI^B4-20^ (**Fig. 4&5**) at an MOI from 0.2 to 0.00002 and 51 days for HAE-ALI^L209^ that was polarized on the large MilliCell™ insert at an MOI 0f 0.2 (**SFig. 3&4**). At an MOI of 0.2 pfu/cell, the infection of HAE-ALI^B4-20^ clearly displayed three peaks at 4, 15, and 31 dpi (**Fig. 4A**), and the infection of HAE-ALI^L209^ displayed 5-6 peaks at 3, 14, 23, 31, and 41 dpi (**SFig. 3A**). There were significant numbers of infected cells (NP+) at 31 and 51 dpi in HAE-ALI^B4-20^ and HAE-ALI^L209^, respectively (**Fig. 5**, MOI=0.2 and **SFig. 4**). Again, the barrier function of the infected HAE-ALI was diminished, as determined by the TEER measurement, begun at 1 dpi (**Fig. 4B&D** and **SFig. 3B**), as well as by the dispersed ZO-1 staining (**Fig. 5A**, MOI=0.2, and **SFig. 4A**). We also consistently observed drastic loss of the cilia (**Fig. 5B**, MOI=0.2, and **SFig. 4B**).

**Figure 4.**
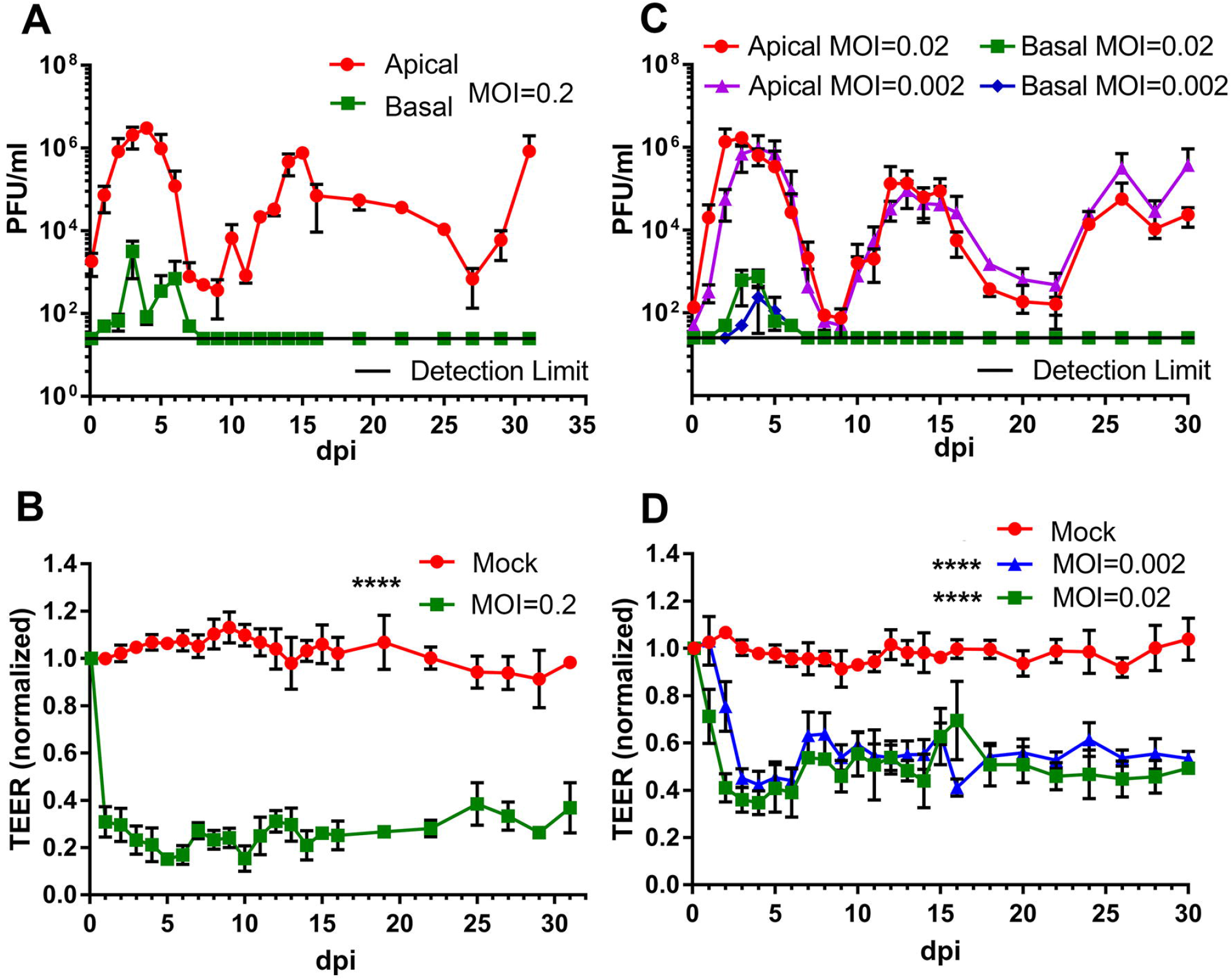
Virus release kinetics and transepithelial electrical resistance (TEER) measurement of HAE-ALI infected with SARS-CoV-2 at various viral loads (multiplicities of infection). **(A&C) Virus release kinetics.** HAE-ALI^B4-20^ cultures were infected with SARS-CoV-2 at an MOI of 0.2 (A), 0.02 and 0.002 (C), respectively, from the apical side. At the indicated days post-infection (dpi), 100 μl of apical washes by incubation of 100 μl of D-PBS in the apical chamber and 100 μl of the basolateral media were taken for plaque assays. Plaque forming units (pfu) were plotted to the DPI. Value represents the mean +/− standard deviations. (**B&D**) **Transepithelial electrical resistance measurement.** The TEER of mock- and SARS-CoV-2-infected HAE-ALI culture were measured using an epithelial Volt-Ohm Meter (Millipore) at the indicated dpi. The TEER values were normalized to the TEER measured on the day of infection, which is set as 1.0. Values represent the mean of relative TEER +/− standard deviations. **** P < 0.0001 by one-way Student t-test.

**Figure 5.**
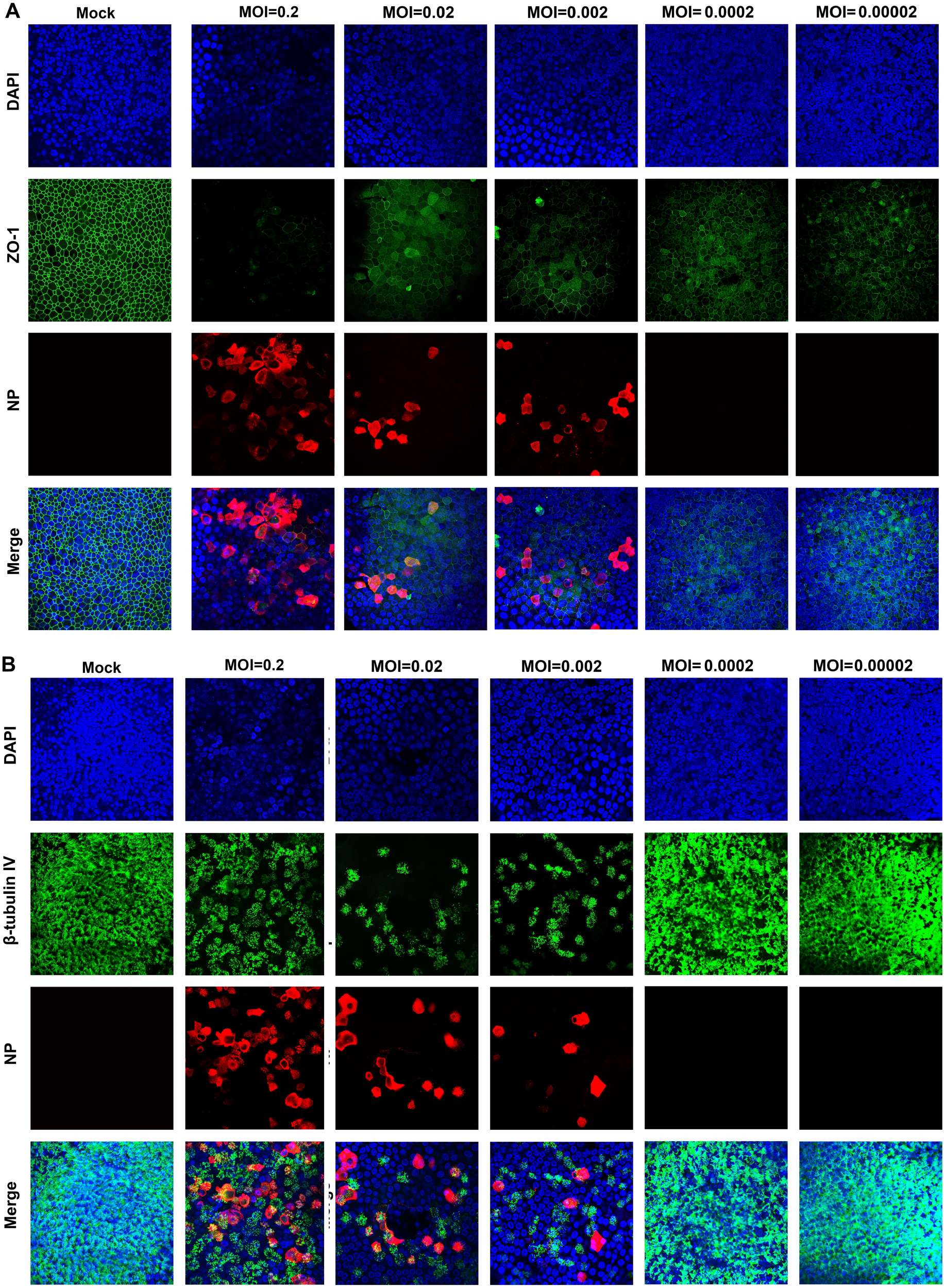
Immunofluorescence analysis of SARS-CoV-2 infected primary bronchial HAE at various viral loads (multiplicities of infection). HAE-ALI^B4-20^ cultures were infected with SARS-CoV-2 at an MOI from 0.2 to 0.00002. At 30 dpi, both virus and mock infected HAE were co-stained with anti-NP and anti-ZO-1 antibodies (**A**), or co-stained with anti-NP and anti-β-tubulin IV antibodies (**B**). Confocal images were taken at a magnification of x 40. Nuclei were stained with DAPI (blue).

Viral shedding to the culture medium in the basolateral chamber across the supportive membrane was also continuously monitored in the experiments. The infectious virions in the medium were detectable in the early time points after the infection was initiated, but at a level of two to three orders of magnitude less than that in the apical washes (**Fig. 4A**, **SFig. 1A**, and **SFig. 3A**), suggesting the that the progeny of the SARS-CoV-2 are predominately released from the apical membrane of the infected HAE. We concluded that the trace of detectable viruses in the basolateral medium must come from the leakage of the apically secreted viruses across the supportive semipermeable membrane due to epithelial damage caused from SARS-CoV-2 infection. Interestingly, no infectious virions were found in the basal medium when the peaks of the viral progeny in the apical washes reappeared at the late time points, even though virus burdens were at similar levels at these peaks (**Fig. 4A&C**, **SFig. 1A**). These observations suggest the regeneration of the destructive mucosal lesions occur during the SARS-CoV-2 infection and that such repair is sufficient to prevent the viral shedding to the basolateral chamber, although the repair did not lead to the recovery of the TEER of the infected HAE (**Fig. 4B&D**). Different observation was from the infection of HAE-ALI^L209^, in which the epithelial cells from donor L209 were cultured and polarized in the large Millicell™ inserts (1.1 cm^2^). Traces of viral shedding in the basal medium were also found at the late time points and their levels changed responding to each peak of the virus replication (**SFig. 3A**). We reasoned that this is due to the inefficient epithelium repair in the infected HAE-ALI^L209^, which was affected by either donor variation or different culture format.

The recurrent peaks of virus progeny released during the course of infection were further displayed in infections of HAE-ALI^B4-20^ at lower MOIs of 0.02 and 0.002 (**Fig. 4C**), which show three almost identical peaks at 3, 13, and 26 dpi over the course of 30 days. Of note, the 3^rd^ peak became obvious at 26 dpi from these low MOI infections, compared to 30 dpi for the higher MOI of 0.2, but the amounts of infectious virions released from those peaks remained at roughly the same level (~1 × 10^6^ pfu/ml). The epithelial damage was indicated by the decrease of TEER beginning at 2 and 3 dpi, respectively (**Fig. 4D**), and also revealed by the dispersed ZO-1 expression and loss of cilia (**Fig. 5**, MOIs=0.02 and 0.002).

We then further carried out the infection at much lower MOIs, 0.0002 and 0.00002, receptively, over a course of 3 weeks. Surprisingly, we found that HAE-ALI^B4-20^ cultures were not infected by SARS-CoV-2, as evidenced by no NP+ cells at 30 dpi. (**Fig. 5**, MOI=0.0002 and 0.00002) and no infectious virions released from the apical side were detectable. To verify this result, we performed infections in HAE-ALI^B9-20^ at MOIs of 0.0002 and 0.00002, which reproduced the same observations of no productive infection in the cultures derived from a different lung donor (**SFig. 5**). At MOIs of 0.0002 and 0.00002, we did not observe an obvious loss of cilia in both infected HAE-ALI^B4-20^ and HAE-ALI^B9-20^ (**Fig. 5B** and **SFig. 5B**); however, we observed a cytoplasmic expression and a weak junction expression of ZO-1 at 30 dpi (**Fig. 5A)** and 21 dpi (**SFig. 5A**) for infected HAE-ALI^B4-20^ and HAE-ALI^B9-20^, respectively. These results demonstrate that a high viral load (at least >100 pfu (~8.2 ×10^4^ vgc) to an epithelium of 0.33 cm^2^, which contains with ~5 ×10^5^ epithelial cells, is necessary to initiate a productive infection.

### Ciliated and goblet cells are permissive to SARS-CoV-2 but not the basal and club cells

We next examined SARS-CoV-2 infection in which the inoculation of a high MOI of 2 was applied to the basolateral side in HAE-ALI^B4-20^. The results showed there were no detectable infectious virions released from both the apical and basolateral sides (**Fig. 6A**). The TEER of the infected HAE displayed no significant changes over the course of 24 days (**Fig. 6B**), which was also evidenced for the well-preserved tight junctions (**Fig. 6C**), as well as the rich cilia expression (**Fig. 6D**). Importantly, NP+ cells were not detected for as long as 23 dpi. Similar results were verified in infection of HAE-ALI^B9-20^ over an infection course of 3 weeks. These results demonstrated SARS-CoV-2 does not infect epithelial cells from the basolateral side.

**Figure 6.**
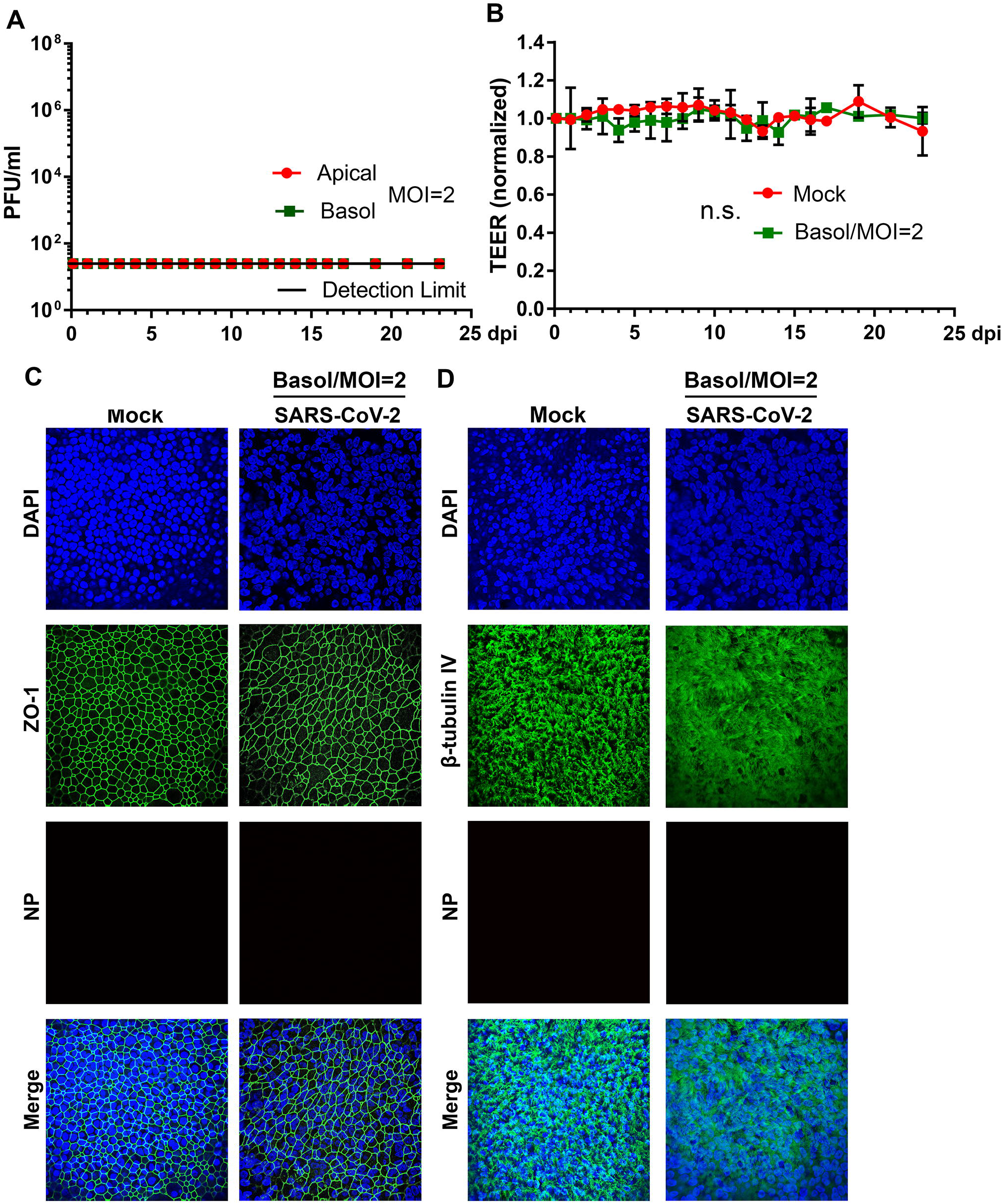
SARS-CoV-2 does not infect HAE-ALI from the basolateral side. **(A) Virus release kinetics.** Both apical washes and basolateral media were collected from SARS-CoV-2 infected HAE-ALI^B4-20^ every day and quantified for virus titers using plaque assays. Plaque forming units (pfu) were plotted to the dpi. Value represents the mean +/− standard deviations. **(B) Transepithelial electrical resistance (TEER) measurement.** The TEER of infected HAE-ALI^B4-20^ cultures were measured using an epithelial Volt-Ohm Meter (Millipore) at the indicated dpi, and were normalized to the TEER measured on the first day, which is set as 1.0. Values represent the mean of the relative TEER +/− standard deviations. n.s. indicates statistically no significance. **(C&D) Immunofluorescence analysis.** Mock- and SARS-CoV-2-infected HAE-ALI^B4-20^ cultures at 23 dpi were co-stained with anti-NP and anti-ZO-1 antibodies (**C**), or co-stained with anti-NP and anti-β-tubulin IV antibodies (**D**). Confocal images were taken at a magnification of x 40. Nuclei were stained with DAPI (blue)

To determine the permissive epithelial cell types in the infected HAE-ALI, we carefully examined the infected cells by immunofluorescent assays using various epithelial cell markers. Cells were dissociated from the supportive membrane of the infected Transwell^®^ inserts and cytospun onto slides for imaging. Co-staining of a specific cell marker and the viral NP expression visualized the cell types permissive for SARS-CoV-2 infection. The results, as the representative images shown in **Fig. 7**, demonstrated that the majority of cell populations in the HAE-ALI were basal cells, which expressed cytokeratin 5 (CKRT5+) ^41^, and ciliated cells with positive anti-β-tubulin IV staining. Consistent with previous imaging results (**Fig. 3**), most of the NP+ cells were also positive with anti-β-tubulin IV staining (**Fig. 7A**), whereas almost all the CKRT5+ basal were negative for anti-NP staining (**Fig. 7C**). Probing secretoglobin family 1A member 1 (SCGB1A1) expression for club cells and mucin 5AC (MUC5AC) expression for goblet cells ^41^ revealed that the secretory cells were less abundant sub-populations in the infected HAE-ALI cultures. While we could not locate any club cells stained positively for NP expression (**Fig. 7D**), we found some NP+/MUC5AC+ goblet cells (**Fig. 7B**). Importantly, we observed that in the infected HAE-ALI, a subset of CYKT5+ basal cells are found associated with the expression of Ki67; however, in the mock-infected HAE-ALI, we did not found the cells co-expressing both Ki67 and CYKT5 (**Fig. 8B**). As Ki67 is a marker of cell proliferation ^42^, this result suggested that SARS-CoV-2 infection activates basal cells towards proliferation.

**Figure 7.**
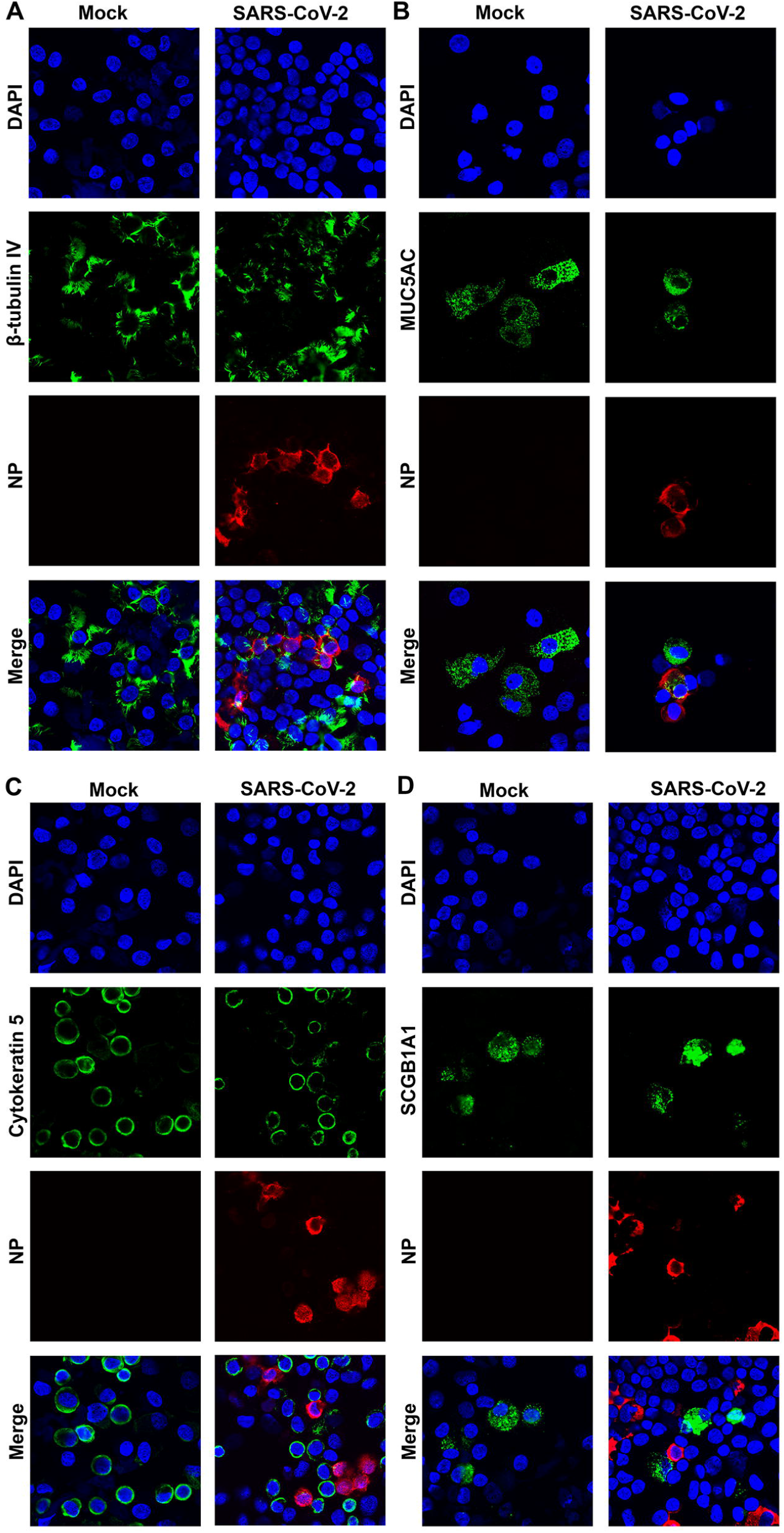
SARS-CoV-2 infects ciliated and goblet epithelial cells but not basal and club cells. Epithelial cells of the mock- and SARS-CoV-2-infected HAE-ALI^B9-20^ cultures at 4 dpi (MOI=0.2) were dissociated from the Transwell^®^ insert and cytospun onto slides. The cells on the slides were fixed, permeabilized, and immunostained with anti-NP and together with anti-β-tubulin IV (**A**), and anti-MUC5AC (**B**), anti-cytokeratin 5 (**C**), and anti-SCGB1A1 (**D**), respectively. Confocal images were taken at a magnification of 63 ×. Nuclei were stained with DAPI (blue).

**Figure 8.**
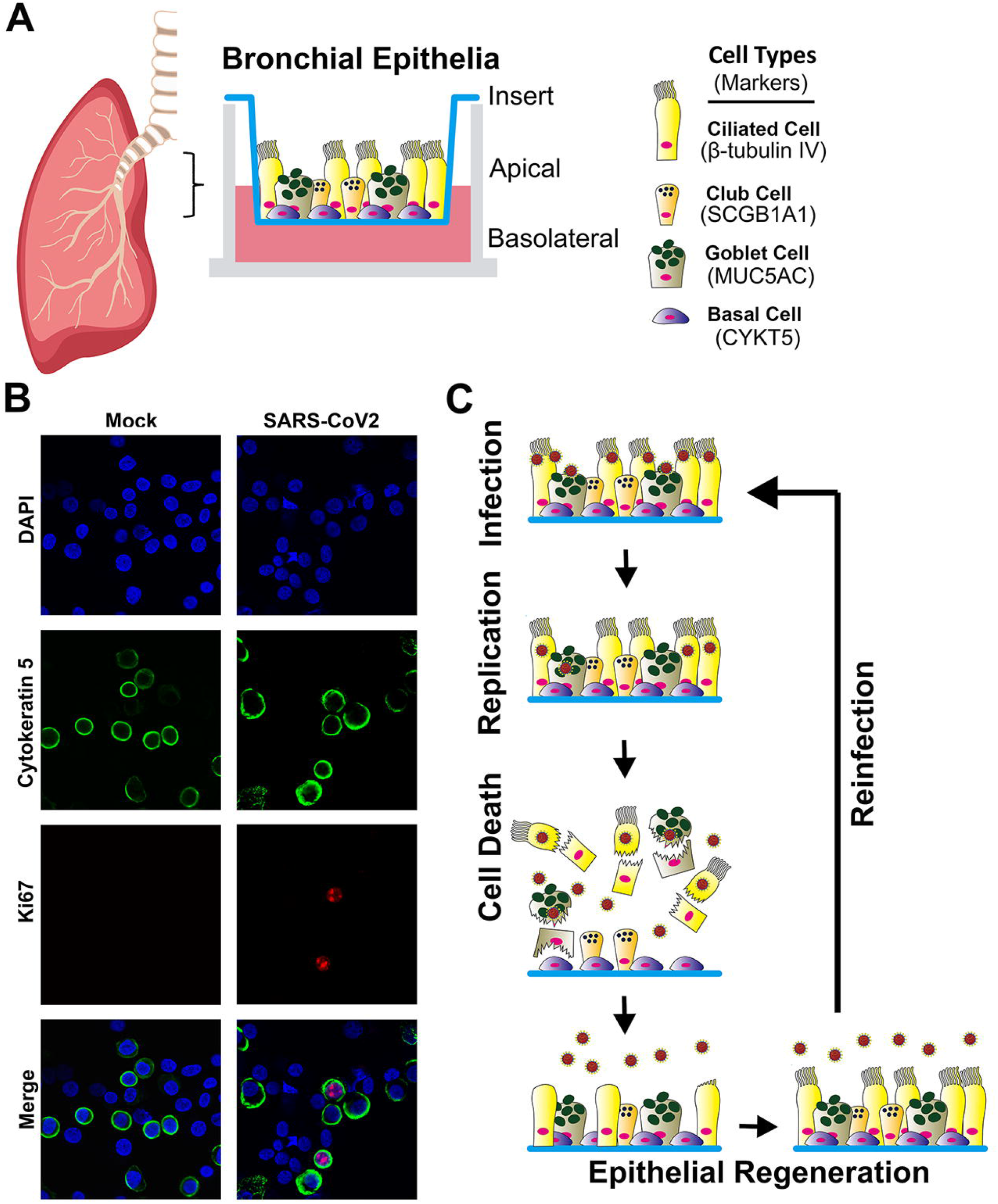
A diagram of HAE-ALI and model of the SARS-CoV-2 recurrent infection in HAE. (**A**) **HAE-ALI model:** Epithelial cells are taken from bronchia of the lung of healthy donors and plated onto Transwell^®^ inserts at an air-liquid interface (ALI) for four weeks. Four major types of the epithelial cells in the well differentiated polarized HAE cultures: basal, ciliated, goblet, and club cells are diagrammed in the Transwell^®^ insert, and their expression makers are indicated. (**B**) **Basal cells in proliferation.** Epithelial cells of the mock- and SARS-CoV-2-infected HAE-ALI^B9-20^ cultures at 9 dpi (MOI=0.2) were dissociated from the Transwell^®^ insert and cytospun onto slides. The cells on the slides were fixed, permeabilized, and immunostained with anti-Ki67 and together with anti-CYKT5. Confocal images were taken at a magnification of 63 ×. Nuclei were stained with DAPI (blue). (**C**) **Model of airway cell regeneration model of SARS-CoV-2 recurrent infections.** SARS-CoV-2 infects apical ciliated and goblet cells, in which it replicates to produce infectious progeny and causes the death of the infected cells. The destructive lesion of epithelium induces basal cell proliferation and differentiation to regenerate ciliated and goblet cells, which are readily infected by SARS-CoV-2 in the next cycle of the recurrent infections.

Taking these lines of evidence together, our results confirmed SARS-CoV-2 mainly infects ciliated cells of HAE, as well as the goblet cells, despite its lower abundance in HAE-ALI cultures. Our study suggests that basal and club cells are not permissive to SARS-CoV-2.

## Discussion

In this study, we modeled SARS-CoV-2 infection in HAE-ALI cultures generated from primary bronchial epithelial cells directly isolated from 4 lungs of independent donors without cell expansion. The infections were conducted at various MOIs, from both apical and basolateral sides, and for a long period from 21-51 days. Our studies demonstrated that the SARS-CoV-2 infection of HAE results in multiple replication peaks of virus progeny at MOIs from 2 to as low as 0.002, although the length of peak emergence time varied from ~7-10 days. The most striking result we obtained is the resistance to SARS-CoV-2 infection at an MOI of 0.0002 (~300 pfu/cm^2^ of epithelium). Our studies also revealed that the basal (CKRT5+) cells and club (SCGB1A1+) cells are not infected, whereas ciliated cells (β-tubulin IV+) and goblet (MUC5AC+) cells are the primary body of the infected cells in the SARS-CoV-2-infected HAE.

Whole-mount immunohistochemistry of SARS-CoV-2 infected HAE cultures revealed that ciliated cells were the cell type predominately infected ^38^. With the single epithelia cell suspension recovered from the infected inserts, we prepared the cytospins to investigate in detail the cell types permissive to the infection. It is very interesting that secretory goblet and club cells behave oppositely to SARS-CoV-2 infection, since both of these two cell types are the surface epithelial cells accessible on the airway lumen ^38^. The permissibility of goblet cells to SARS-CoV-2 infection was previously reported by immunochemistry staining of biopsy airway section specimens from patients of COVID-19 ^36^, as well as being supported by a recent in vitro modeling of HAE ^34^. Club cells express the viral receptor ACE2, but at a less abundant level than goblet cells and ciliated cells ^43^. The finding of club cell’s failure to establish productive SARS-CoV-2 infection suggests a possible lack of the expression of the entry essential serine protease TMPRSS-2 in this cell type ^44, 45^. Of note, club cells are non-ciliated epithelial cells and they are restricted to just the small airways in humans ^46^. While club cells are recognized as secretory cells of major importance to the epithelium, they also act as a stem cell, multiplying and differentiating into ciliated cells and goblet cells to regenerate the bronchiolar epithelium ^47^. In mouse trachea, the club cells exist as a transiently amplifying population, and their capacity for self-renewal and multilineage differentiation is enhanced after injury ^48^. Taking together with the fact that the basal cells are not permissive to SARS-CoV-2 infection but the basal cell proliferation was activated after SARS-CoV-2 infection (**Fig. 8B**), the basal and possibly also the club cell types should play an important role in repairing the epithelium lesions caused by viral infection.

Our observation that SARS-CoV-2 was unable to infect epithelial cells from the basolateral side supports that the viral entry receptor ACE2 is polarly expressed at the apical side ^30, 31^. We believe the finding that CKRT5+ basal cells are largely not infected by SARS-CoV-2 is important to understand the course of SARS-CoV-2 infection in HAE. The airway basal cells are the epithelial cell type not presenting on the surface of the airway lumen, thus, they are not usually accessible to the virus on the apical side. However, when the infection commences and the epithelial damage occurs, the destructive mucosal lesions (and the death of the infected ciliated and goblet cells) would allow the virus to get access the basal cells (**Fig. 8C**). Indeed, the detectable virus shedding to the basolateral chamber (**Fig. 4A&C**, **SFig. 1A** and **SFig. 3A**) indicates a possible window to expose the basal cells to SARS-CoV-2. Notably, these time points also represent the peaks of the release of virus progeny. However, none of the CKRT5+ cells that was also NP-positive was found from the cytospins prepared from SARS-COV-2 infected HAE at 9 dpi when the infection appeared at the lowest level. We hypothesize that the non-permissive nature of basal cells to SARS-CoV-2 is due to the negligible expression of ACE2 ^49^ or TMPRESS-2 ^43^.

The epithelial cell lining of the airways provides an efficient barrier against microbes and aggressive molecules through interdependent functions, including mechanical clearance of the mucus executed by movements of the cilia, a cellular barrier function by means of intercellular epithelial junctions formed by a set of tight junction associated proteins such as ZO1, and homeostasis of ion transport ^40^. At the airway epithelial cellular level, the tight junction– associated proteins, such as ZO1, occludin, and claudins, play a central part in the epithelial cytoprotection by maintaining a physical selective barrier between external and internal environments. The tight junction proteins are highly labile structures whose formation and structure may be very rapidly altered after airway injury, for example, airway inflammation. Proinflammatory cytokines have a drastic effect on tight junction expression and barrier functions, which significantly alter the epithelial barrier permeability ^50–52^. SARS-CoV-2 infection distorted the ZO-1 expression, and thereafter causes barrier dysfunction (TEER decrease). The infection not only alters the ZO-1 expression of infected (NP1+ cells) but also uninfected cells (NP-cells) (**Fig. 3**). This is also true for the cilia loss. We believe that SARS-CoV-2 infection produces inflammatory cytokines as an innate immunity response upon virus infection ^53^, which either disturbs ZO-1 and tubulin expression or alters their structures. The innate immunity response also limits virus infection at the front line. In fact, SARS-CoV-2 requires a high viral load (>300 pfu/cm^2^ of HAE) to initiate a productive infection (**Fig. 4**). Of note, the infectious titer (pfu) was determined by plaque assay in Vero-E6 cells, which are interferon-deficient ^54^. We determined that 1 pfu of SARS-CoV-2 in Vero-E6 cells has a particle (viral genome copy) number of 820, suggesting that a load of 2.46 × 10^5^ particles is required to productively infect 1 cm^2^ of the airway epithelium, which is much higher than the small DNA virus parvovirus human bocavirus 1 (HBoV1) we studied ^55^. HBoV1 can infect HAE at an MOI of as low as 0.001 genome copies per cell, which equals 1.5 × 10^3^ particles per 1 cm^2^ of the airway epithelium.

Epithelial regeneration involves migration of the basal cells that neighbor the acute injured area (e.g., virus-infected area), active dividing and squamous metaplasia, rapid redifferentiation to preciliated cells, and finally ciliogenesis towards a complete pseudostratified mucociliary epithelium ^56^. Airway epithelium repair is critical for the maintenance of the barrier function and the limitation of airway hyperreactivity. In a biopsy study of fresh tracheas and lungs from five deceased COVID-19 patients, it was found that the epithelium was severely damaged in some parts of the trachea, and extensive basal cell proliferation was observed in the trachea, where ciliated cells were damaged, as well as in the intrapulmonary airways ^37^. These data support our conclusion that basal cells are not permissive to SARS-CoV-2. As a response to these previous findings, our study observed that a subset of proliferating basal cells in the SARS-CoV-2 infected HAE-ALI, but not in the mock infected HAE-ALI (**Fig. 8B**). Thus, we hypothesize that SARS-CoV-2 infection induces basal cell proliferation, which accounts for the observed long-lasting infections with recurrent peaks of viral replication, which warrants future investigation.

Overall, we propose a model of SARS-CoV-2-infection of HAE (**Fig. 8C**): SARS-CoV-2 selectively infects ciliated and goblet cells on the surface of the airway lumen (the apical side of HAE). Upon invading these cells, SARS-CoV-2 replicates and produces infectious virions, and eventually leads to the cell death and epithelial damage. Upon the destructive lesions, airway epithelium has the capacity to progressively repair and regenerate itself. Thus, basal cells (possibly also including club cells) proliferate and differentiate to ciliated cells or goblet cells to fill up the areas that have lost ciliated or goblet cells. Then, the virus released from the last round of infection infects newly regenerated ciliated or goblet cells (repaired epithelia), followed by the second round of active replication and virus production. Therefore, the airway epithelial regeneration confers a persistent, cyclically peaked infection of SARS-CoV-2 in human epithelia.

## Methods

### Ethics statement

Primary human bronchial epithelial cells were isolated from the lungs of healthy human donors by the Cells and Tissue Core of the Center for Gene Therapy, University of Iowa and the Department of Internal Medicine, University of Kansas Medical Center with the approvals of the Institutional Review Board (IRB) of the University of Iowa and University of Kansas Medical Center, respectively.

### Cell line and virus

#### Cell line

Vero-E6 cells (ATCC^®^ CRL-1586™) were cultured in Dulbecco’s modified Eagle’s medium (DMEM) (HyClone, catalog no. SH30022.01; GE Healthcare Life Sciences, Logan, UT) supplemented with 10% fetal bovine serum (FBS; catalog no. F0926, Sigma, St. Louis, MO) at 37°C under a 5% CO_2_ atmosphere.

#### Virus

SARS-CoV-2 (NR-52281), isolate USA-WA1/2020, was obtained from BEI Resources, NIAID, NIH. The viruses were propagated in Vero-E6 cells once, titrated by plaque assay, aliquoted in Dulbecco’s phosphate buffered saline, pH7.4; D-PBS, and stored at −80°C in the BSL3 Lab (Hemenway 4037) of the University of Kansas Medical Center. A biosafety protocol to work on SARS-CoV-2 in the BSL3 Lab was approved by the Institutional Biosafety Committee (IBC) of the University of Kansas Medical Center. The virus titers of the wild-type SARS-CoV-2 was 1.0 × 10^7^ pfu/ml, which equals a physical titer of 8.2 × 10^9^ viral genome copies (vgc)/ml determined by reverse transcription-quantitative PCR (RT-qPCR).

### Primary airway epithelium (HAE) cultured at an air-liquid interface (ALI; HAE-ALI) (Fig. 8A)

The primary HAE-ALI cultures HAE-ALI^B3-20^, HAE-ALI^B4-20^, and HAE-ALI^B9-20^ were provided by the Cells and Tissue Core of the Center for Gene Therapy, University of Iowa ^18, 57–59^. These polarized HAE-ALI cultures were derived from three independent donors. The freshly isolated human bronchial epithelial cells from the donor tissues were seeded onto collagen-coated, semipermeable polycarbonate membrane inserts (0.33 cm^2^, 0.4 μm pore size, Costar Transwell^®^, Cat #3413, Corning, New York), and grown at an ALI as previously described ^60^. The cultures were maintained in USG medium containing 2% Ultroser G (USG) serum substitute (Pall BioSepra, France). After 3-4 weeks of culture at an ALI, the polarized culture was fully differentiated. The polarity of the HAE was determined for the transepithelial electrical resistance (TEER) using an epithelial Volt-Ohm Meter (Millipore). A value of 1,000 Ω.cm^2^ or higher was chosen for SARS-CoV-2 infection as we previously used for HBoV1 infection ^52, 61^. HAE-ALI^L209^ cultures on 1.1 cm^2^ Millicell™-PCF (Millipore, Billerica, MA) were provided by Dr. Matthias Salathe, which were generated following a published method ^62^ using primary airway bronchial epithelial cells isolated from the lung of a donor (L209).

### Virus infection, sample collection and titration

#### Virus infection

For apical infection, well-differentiated primary HAE-ALI in Transwell^®^ inserts (0.33 cm^2^; Costar) or in Millicell™ inserts (1.1 cm^2^; Millipore) were inoculated with 100 μl or 300 μl of SARS-CoV-2 at various MOIs, as indicated in each figure legend, applied to the apical chamber. For basolateral infection, HAE-ALI cultures in Transwell^®^ inserts were inoculated with SARS-CoV-2 diluted in 500 μl of USG medium added to the basolateral chamber. The infected HAE-ALI cultures were incubated at 37°C 5% CO_2_ for 1 h followed by aspiration of the virus from the apical or basolateral chamber and washing of the cells with D-PBS three times (the last wash was saved and used for plaque assay, which was presented as the virus residue right after infection at the day 0 post infection (0 dpi)). The HAE-ALI cultures were then further cultured at 37°C and 5% CO_2_.

#### Viral sample collection

Viral samples were collected from both the apical wash of the epithelium surface and the culture medium in basolateral chamber at multiple time points. In brief, 100 μl (or 300 μl) of D-PBS was added to the apical chamber for a short incubation of 30 min at 37ºC and 5% CO_2_. Thereafter, this apical wash was recovered carefully from the apical chamber without disturbing the culture. To quantitate the viruses released from the basal membrane to the culture medium, 100 μl of medium were collected from each basolateral chamber. The infectious titers in the collected samples were determined by plaque assays in Vero-E6 cells.

#### Plaque assays

Vero-E6 cells were seeded in 24-well plates at a density of ~0.5 ×10^6^ cells and grown to confluence the second day. Virus (apical washes or basolateral media) were serially diluted at 10-fold in D-PBS. 200 μl of the diluent were added to each well and incubated for 1 h on a rocking rotator. After removing the virus diluent, ~0.5 ml of overlay media (1% methylcellulose (Sigma, M0387) in DMEM with 5% FBS) were added to each well. The plates were incubated at 37°C under 5% CO_2_ for 4 days. After removing the methylcellulose overlays, cells were fixed using the 10% formaldehyde solution for 30 min and stained with 1% crystal violet solution followed by extensive washing using distilled water. Plaques in each well were manually counted and multiplied by the dilution factor to determine the virus titer at the unit of pfu/ml.

#### RT-qPCR

To eliminate free viral RNA in the samples, 25 units of Benzonase (Sigma) was added to 100 μl of the virus samples for 30 min ^63^. The nuclease-treated samples were used for viral RNA extraction using the Viral RNA extraction kit (Quick-RNA Viral Kit, R1035, Zymo Research) following the manufacturer’s instructions. M-MLV reverse transcriptase (#M368A, Promega) was used to reverse transcribe viral RNA with the reverse PCR primer according to the manufacturer’s instructions. 2.5 μl of the cDNA was quantified by TaqMan qPCR in a reaction of 25 μl to determine the number of viral genome copies (vgc) using the CDC 2019-nCoV_N1 set of primers and probe, which were synthesized at IDT (Coralville, Iowa). The plasmid pcDNA6B-(SARS-CoV-2)N, which contains the SARS-CoV-2 NP gene (nt 998-2,244), was used as reference control (1 vgc = 7 × 10^−12^ μg) to establish a standard curve for absolute quantification on an Applied Biosystems 7500 Fast system (Foster City, CA).

### Immunofluorescent confocal microscopy

#### Immunofluorescent assay

For analysis of the SARS-CoV-2 infection in the HAE grown on the supportive membrane of the Transwell^®^ inserts, we cut off the membrane from the inserts and fixed them with 4% paraformaldehyde in PBS at 4°C overnight. The fixed membrane was washed in PBS for 5 mins three times and then split into 4 (for 0.33 cm^2^ membrane) or 8 (for 1.01 cm^2^ membrane) pieces for whole-mount immunostaining. For cell marker analysis, we dissociated the cells off the supportive membrane of the Transwell^®^ inserts by incubation with Accutase (Innovative Cell Technologies, Inc. San Diego, CA). After incubation for 1h at 37°C, cells were completely detached from the membrane and well separated. Cells were collected and then cytocentrifuged at 18,000 rom for 3 min onto slides using a Shandon Cytospin 3 cytocentrifuge. After cytospun, the slides were fixed overnight in 4% paraformaldehyde at 4°C.

The fixed HAE or dissociated cells were permeabilized with 0.2% Triton X-100 for 15 min at room temperature. Then, the slide was incubated with primary antibody in PBS with 2% FBS for 1 h at 37ºC. After washing, the membrane was incubated with fluorescein isothiocyanate-and rhodamine-conjugated secondary antibodies, followed by staining of the nuclei with DAPI (4’,6-diamidino-2-phenylindole).

#### Confocal microscopy

The cells were then visualized using a Leica TCS SPE confocal microscope at the Confocal Core Facility of the University of Kansas Medical Center. Images were processed with the Leica Application Suite X software.

### Antibodies used

Primary antibodies used were rabbit monoclonal anti-SARS-CoV-2 nucleocapsid (NP) (Clone 001, #40143-R001, SinoBiological US, Wayne, PA) at a dilution of 1:25, mouse monoclonal anti-β-tubulin IV antibody (clone ONS.1A6, #T7941, MilliporeSigma, St Louis, MO) at 1:100, mouse anti-ZO-1 (Clone 1/ZO-1, #610966, BD Bioscience, San Jose, CA) at 1:100, rabbit anti-Ki67 (Clone SP6, ab1666, Abcam, Cambridge, MA7) at 1:50. Mouse anti-MUC5AC (Santa Cruz Biotechnology, #sc-33667; used at 1:10), mouse anti-cytokeratin k5 (ThermoFisher Invitrogen, #MA5-12596; used at 1:50), and rat anti-SCGB1A1 (ThermoFisher Invitrogen, #MAB4218; used at 1:50) were used to mark epithelial cell types.

### Transepithelial electrical resistance (TEER)

100 μl of D-PBS was added to the apical chamber to determine the TEER using a Millicell ERS-2 Voltohmmeter (MilliporeSigma, Burlington, MA) following a previously used method ^60^.

### Statistics

Virus released kinetics were determined with the means and standard deviations obtained from at least three independent HAE-ALI^B3-20, B3-20, and B9-20^ and from duplicated HAE-ALI^L209^ by using GraphPad Prism version 8.0. Error bars represent means and standard deviations (SD). Statistical significance (P value) was determined by using unpaired (Student) t-test for comparison of two groups.

## Acknowledgments

We are grateful to Drs. Matthias Salathe and Michael Kim and Nathalie Baumlin in the Department of Internal Medicine for providing HAE-ALI^L209^ cultures and helpful discussions during the study. We thank the Cells and Tissue Core of Center for Gene Therapy, the University of Iowa (DK054759) for providing the primary cell cultures. The following reagent was deposited by the Centers for Disease Control and Prevention and obtained through BEI Resources, NIAID, NIH: SARS-Related Coronavirus 2, Isolate USA-WA1/2020, NR-52281. We are indebted to Dr. Pei-Hui Wang at Shangdong University, China for the gifted plasmid pcDNA6B-(SARS-CoV-2)N.

This study was supported by PHS grants AI150877, AI139572, and AI151542 from the National Institute of Allergy and Infectious Diseases. This study was also supported by grant YAN19XX0 from the Cystic Fibrosis Foundation.

## Author contributions

S.H., J.Q., Z.Y. designed the infection experiments with assistance from K.N.; S.H. performed plaque assays, RT-qPCR, immunofluorescent assays, and confocal image processing with assistance from J.Q. and K.N; J.Q. detected transepithelial electrical resistance (TEER); Z.Y and K.V. generated and maintained HAE-ALI cultures; C.A.K. maintained HAE-ALI cultures; S.H. conducted data and statistical analysis; Z.Y. and J.Q. wrote the paper with assistance from S.H.; Z.Y. and J.Q. supervised the project.

## Competing interests

The authors declare no conflict of interest.

## Notes

### Competing Interest Statement

The authors have declared no competing interest.

