## Supplemental Figures for "Long Period Modeling SARS-CoV-2 Infection of in Vitro Cultured Polarized Human Airway Epithelium"

Hao, S. et al

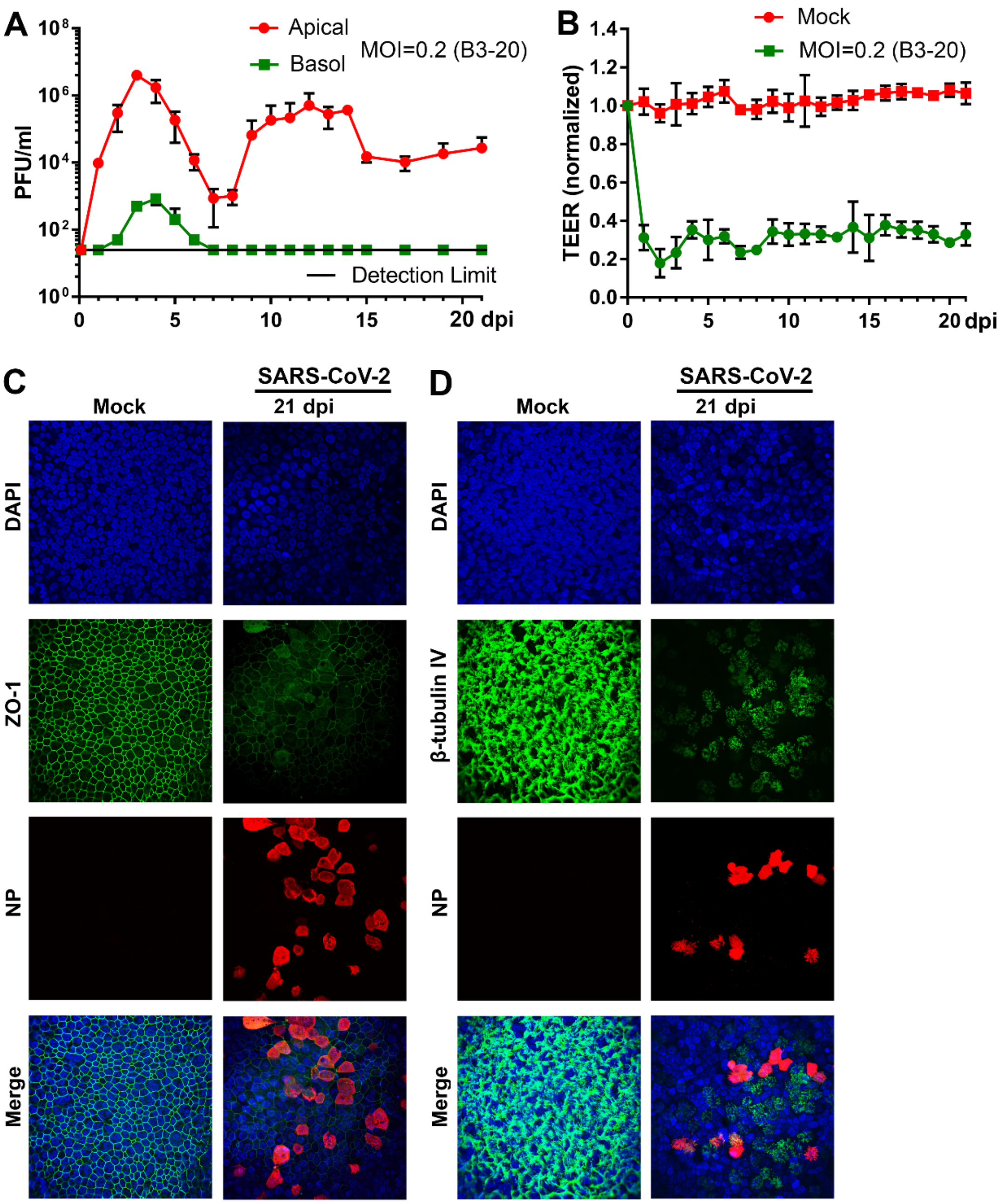


#### Suppl. Figure 1

**Supplementary Figure 1. Virus release kinetics and transepithelial electrical resistance measurement of HAE-ALI$^{B3-20}$ infected with SARS-CoV-2 at MOI of 0.2.**

**(A) Virus release kinetics.** The primary HAE$^{B3-20}$ cultures were infected with SARS-CoV-2 at an MOI of 0.2 from the apical side. At the indicated days p.i. (dpi), 100 µl of apical washes by incubation of 100 µl of D-PBS in the apical chamber and 100 µl of the basolateral media were taken for plaque assays. Plaque forming units (pfu) were plotted to the dpi. Value represents the mean +/- standard deviations. (**B**) **Change of transepithelial electrical resistance (TEER) over the infection period.** The TEER of mock- and SARS-CoV-2-infected HAE$^{B3-20}$ cultures were measured using an epithelial Volt-Ohm Meter (Millipore) at the indicated dpi and were normalized to the TEER measured at the first day, which is set as 1.0. Values represent the mean of the relative TEER +/- standard deviations. **** P< 0.0001. (**C&D**) **Immunofluorescence analysis.** Mock- and SARS-CoV-2-infected HAE-ALI$^{B3-20}$ cultures at 21 dpi were co-stained with anti-NP and anti-ZO-1 antibodies (**C**), or co-stained with anti-NP and anti-β-tubulin IV antibodies (**D**). Confocal images were taken at a magnification of x 40. Nuclei were stained with DAPI (blue).

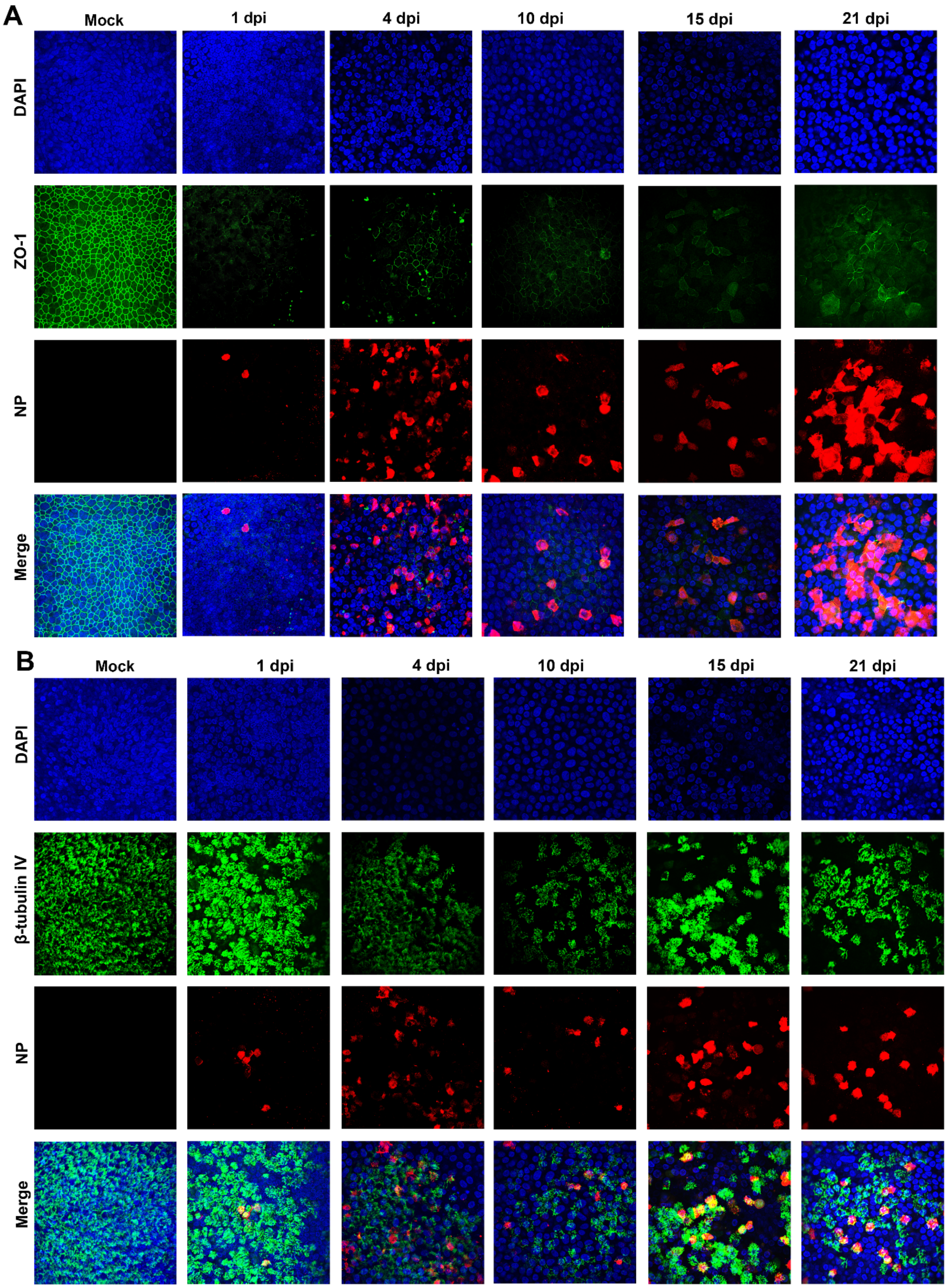


**Suppl. Figure 2**

**Supplementary Figure 2. Immunofluorescence analysis of SARS-CoV-2 infected HAE-ALI$^{B9\text{-}20}$ at an MOI of 2 over a time course of 21 days.**

Mock- and SARS-CoV-2-infected HAE-ALI$^{B9\text{-}20}$ cultures at the indicated days p.i. (dpi) were co-stained with anti-NP and anti-ZO-1 antibodies (**A**), or co-stained with anti-NP and anti-β-tubulin IV antibodies (**B**). Confocal images were taken at a magnification of x 40. Nuclei were stained with DAPI (blue).

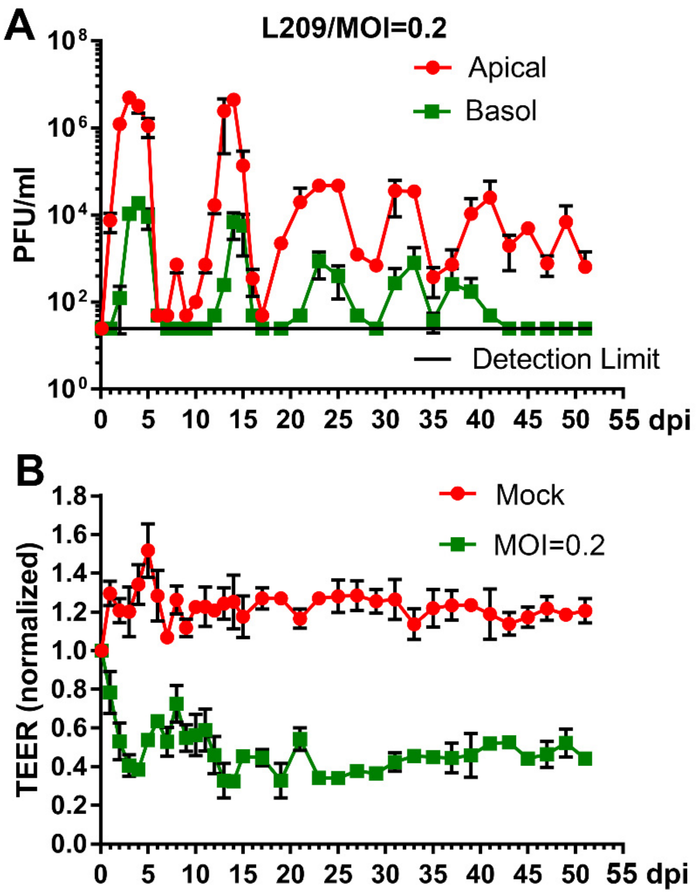


### Suppl. Figure 3

**Supplementary Figure 3. Virus release kinetics and transepithelial electrical resistance measurement of HAE-ALI$^{L209}$ infected with SARS-CoV-2 at MOI of 0.2.**

**(A) Virus release kinetics.** The primary HAE-ALI$^{L209}$ cultures were infected with SARS-CoV-2 at an MOI of 0.2 from the apical side. At the indicated days post-infection (dpi), 300 µl of apical washes by incubation of 300 µl of D-PBS in the apical chamber and 300 µl of the basolateral media were taken for plaque assays. Plaque forming units (pfu) were plotted to the dpi. Value represents the mean +/- standard deviations. (**B**) **TEER measurement.** The TEER of mock- and SARS-CoV-2-infected primary HAE-ALI$^{L209}$ cultures were measured using an epithelial Volt-Ohm Meter (Millipore) at the indicated dpi and were normalized to the TEER measured on the first day, which is set as 1.0. Values represent the mean of the relative TEER +/- standard deviations.

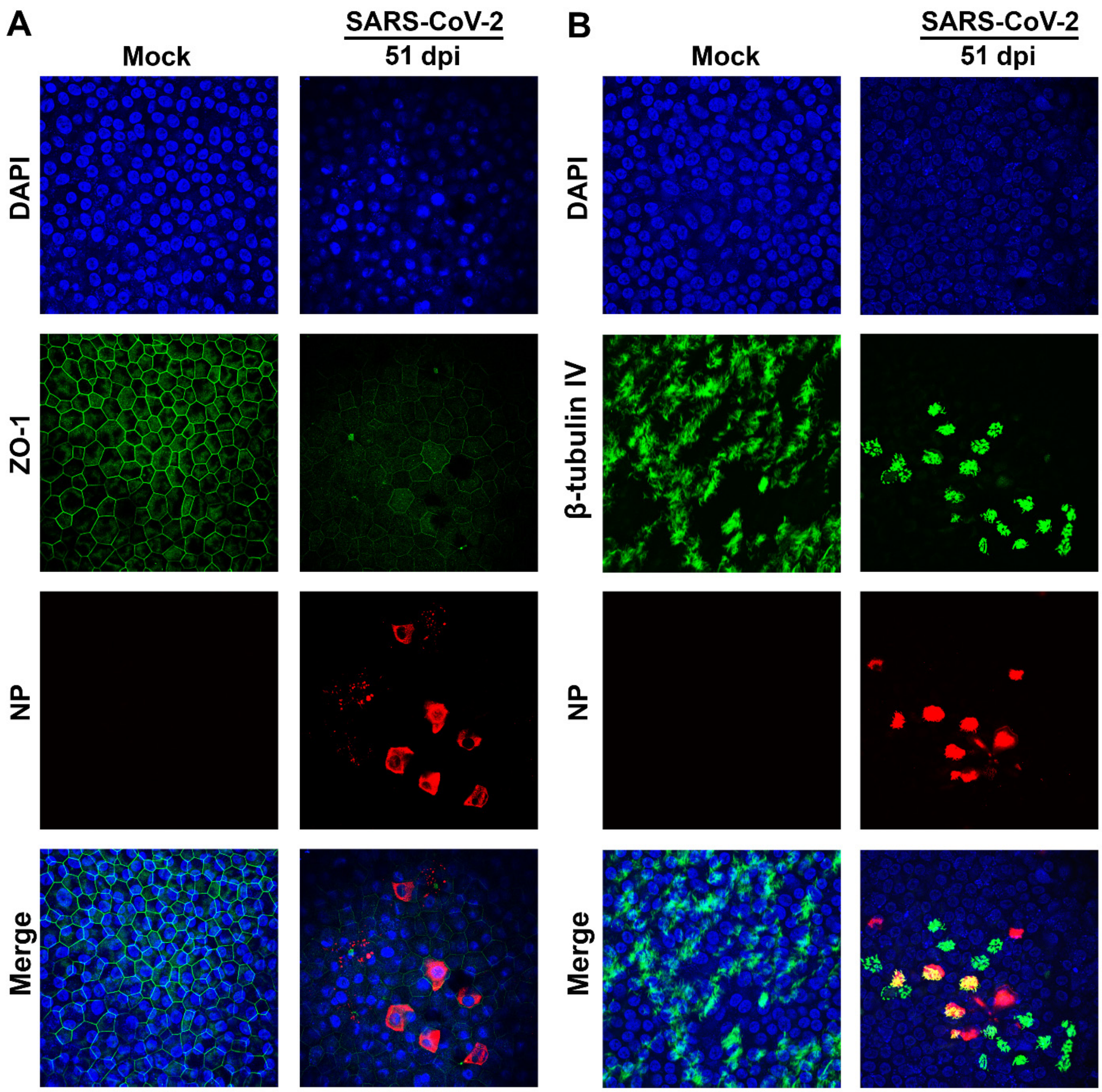


**Suppl. Figure 4**

**Supplementary Figure 4. Immunofluorescence analysis of SARS-CoV-2 infected HAE-ALI$^{L209}$ at an MOI of 0.2.**

Mock- and SARS-CoV-2-infected HAE-ALI$^{L209}$ cultures at 51 dpi were co-stained with anti-NP and anti-ZO-1 antibodies (**A**), or co-stained with anti-NP and anti-β-tubulin IV antibodies (**B**). Confocal images were taken at a magnification of x 40. Nuclei were stained with DAPI (blue).

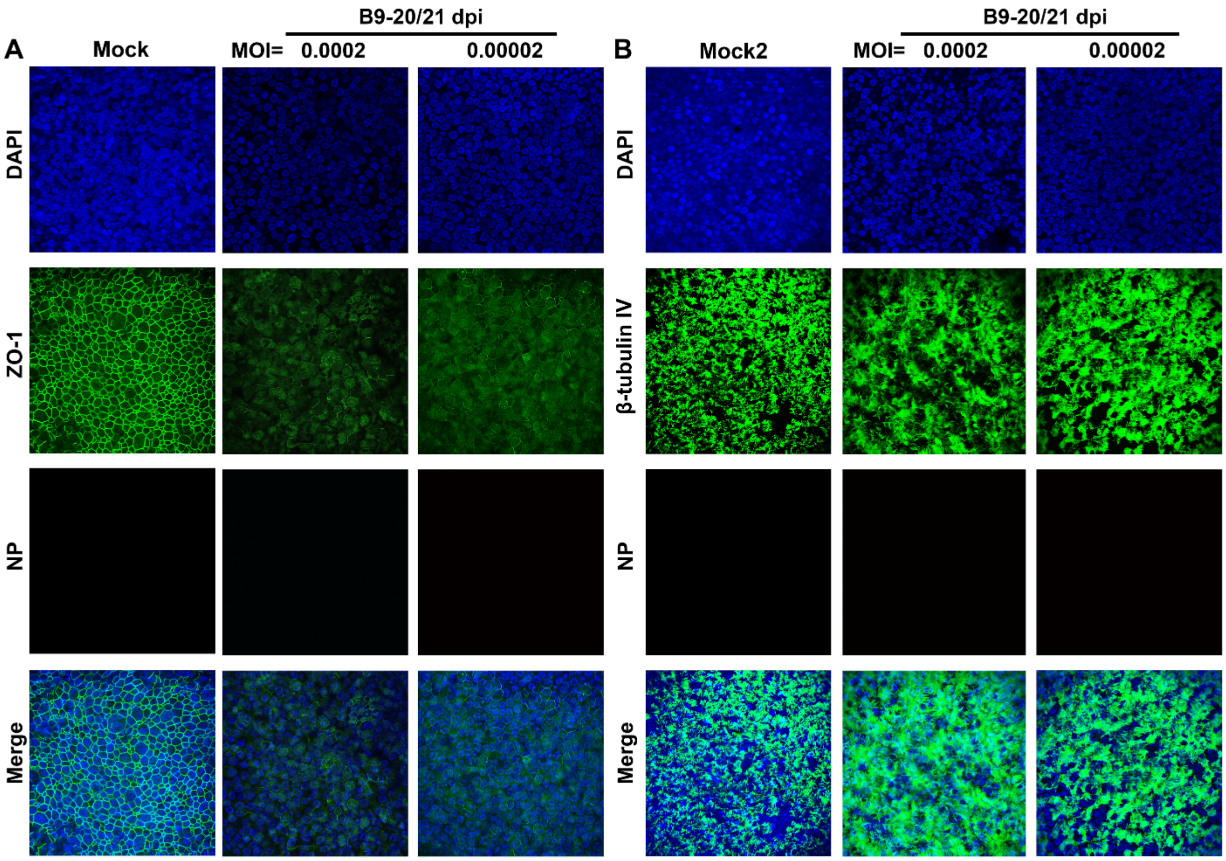


**Suppl. Figure 5**

**Supplementary Figure 5. Immunofluorescence analysis of SARS-CoV-2 infected HAE-ALI$^{B9\text{-}20}$ at MOIs of 0.0002 and 0.00002, respectively.**

Mock- and SARS-CoV-2-infected HAE-ALI$^{B9\text{-}20}$ cultures at 21 dpi were co-stained with anti-NP and anti-ZO-1 antibodies (**A**), or co-stained with anti-NP and anti-β-tubulin IV antibodies (**B**). Confocal images were taken at a magnification of x 40. Nuclei were stained with DAPI (blue).
